# Growth Rates and Metabolic Traits Differ by Diarrheal Manifestation in *Campylobacter jejuni* Strains

**DOI:** 10.1101/2025.04.24.650550

**Authors:** Jennifer M. Bosquez, Craig T. Parker, Ben Pascoe, Kerry K. Cooper

## Abstract

**Introduction:** *Campylobacter jejuni* is the leading cause of bacterial gastroenteritis worldwide. Infections with *C. jejuni* can result in two different diarrheal manifestations in humans: watery diarrhea or bloody/inflammatory diarrhea.

**Hypothesis/Gap Statement:** Currently, little is known about *C. jejuni* and/or host factors associated with elicitation of these two distinct diarrheal manifestations. We hypothesize that these factors may include growth and metabolic trait differences between *C. jejuni* strains associated with watery diarrhea and bloody/inflammatory diarrhea.

**Aim:** Using *C. jejuni* strains with a defined diarrheal manifestation in the neonatal piglet model, we aimed to assess differences in temperature dependent growth rates, motility, biofilm production, and carbon utilization between diarrheal manifestation groups.

**Methodology:** Strains were initially assessed for 192 different carbon sources using phenotypic microarrays followed by specific carbon utilization, growth, motility, and biofilm assays at 37°C and/or 42°C.

**Results:** We found that at 37°C, watery diarrhea associated *C. jejuni* strains grew significantly faster compared to bloody/inflammatory diarrhea associated *C. jejuni* strains. However, there was no significant growth difference at 42°C between the groups, due to bloody/inflammatory diarrhea associated strains growing faster at 42°C compared to 37°C. Additionally at 37°C, we found that L-fucose utilization was significantly higher among watery diarrhea associated strains, while L-glutamine utilization was significantly higher among bloody/inflammatory diarrhea associated strains.

**Conclusion:** The results indicate there are distinct metabolic adaptations between watery and/or bloody/inflammatory diarrhea associated *C. jejuni* strains particularly at 37°C, which may be one of the factors associated with differing diarrheal manifestations.

## Introduction

*Campylobacter jejuni* is a major cause of bacterial gastroenteritis worldwide, responsible for an estimated 550 million cases annually [1]. Infected individuals typically present within 2-5 days of exposure with symptoms including abdominal cramping, nausea, fever, and diarrhea, which can be either watery or bloody/inflammatory in nature [2–4]. Although these distinct diarrheal manifestations have been recognized for decades [5, 6], little is known about the bacterial phenotypic traits that might contribute to these different clinical outcomes.

Watery diarrhea is characterized by loose, non-bloody stools, while bloody/inflammatory diarrhea includes the presence of blood in the stool and/or inflammation of the ileum or colon [4, 7]. Beyond the acute symptoms, campylobacteriosis is associated with several post-infectious sequelae, including Guillain-Barré syndrome and post-infectious irritable bowel syndrome (PI-IBS) [8, 9]. Previous studies suggest that watery diarrhea is linked to secretory mechanisms and enterotoxin activity [10–14], while bloody diarrhea is associated with increased invasiveness [15–17] and cytotoxin production [18–20]. However, fundamental bacterial traits involved in colonization and persistence, such as growth rate, carbon source utilization, motility, and biofilm formation, have not been systematically examined in the context of diarrheal manifestation.

*C. jejuni* exhibits substantial phenotypic diversity, with variation observed even in strains sharing the same single sequence type (ST) [21, 22], lipooligosaccharide (LOS) structure [23, 24], biofilm formation capacity, aerotolerance [25], antimicrobial resistance, immune response modulation [26], and/or association with PI-IBS [27]. Agricultural and laboratory strains demonstrate further variability due to genetic drift [28] and phase-variation [29] following multiple passages [30–34]. Unlike many enteric bacteria, *C. jejuni* lacks pathways necessary for the metabolism of common intestinal carbohydrates such as glucose and galactose [35]. Despite its relatively small genome and fewer identified metabolic pathways compared to other enteric pathogens like *Salmonella, C. jejuni* demonstrates notable metabolic flexibility, utilizing amino acids or citric acid cycle intermediates as energy sources [31, 36, 37].

This metabolic heterogeneity includes the ability of some strains to metabolize glutamine and asparagine [38]. The amino acid L-glutamine can serve as a primary energy source in *C. jejuni* strains encoding a gamma-glutamyl transpeptidase (GGT) [39], with GGT-positive strains, such as those in the ST-45 clonal complex (ST-45 CC) [40], showing enhanced colonization in murine models. Other strains may compensate for the lack of GGT by through alternative metabolic pathways [38, 41]. Certain strains, particularly those of ST-22 CC [40] harbor a genomic island for L-fucose metabolism - a sugar found in intestinal mucin glycoproteins [42, 43]. L-fucose metabolism confers a competitive advantage during colonization in some hosts, though its role in human pathogenesis remains unclear [42, 44]. These metabolic differences may contribute to variations in colonization potential, virulence, and clinical manifestation.

Despite these known differences, colonization factors such as growth, motility, biofilm formation, and carbon utilization have not been systematically studied for their potential roles in influencing diarrheal outcomes of *C. jejuni* infections. Addressing this knowledge gap is essential for better understanding the pathogenesis of this important pathogen. In this study, we investigated whether growth characteristics and metabolic capacity differ between strains of *C. jejuni* associated with distinct diarrheal manifestations, and whether these differences reflect potential host adaptation or zoonotic traits. Using a panel of ten well-characterized *C. jejuni* strains shown to cause either watery or bloody/inflammatory diarrhea in the neonatal piglet model - a system that reliably recapitulates human disease phenotypes [45, 46], we compared generation times, carbon source utilization (L-fucose, L-glutamine, and mucin), motility, and biofilm formation across temperatures representative of human (37°C) and chicken (42°C) hosts.

## Materials and Methods

### Bacterial Culturing and Growth

All strains used in this study, their source, year of isolation, and associated diarrheal manifestation in the neonatal piglet model are listed in **Table 1**. The strains selected for this study were chosen based on their documented diarrheal manifestations in neonatal piglets, offering a more controlled way to investigate the mechanisms behind the observed clinical variability. All watery diarrhea associated strains were sourced from chickens (*Gallus gallus*) while two strains associated with bloody/inflammatory diarrhea originated from chickens and the other three were isolated from human clinical cases. In addition to the *in vivo* work that has been performed with these strains, all have been characterized either genotypically or phenotypically to varying degrees in previous studies, with strains NCTC11168, 81-176 and M129 the most well characterized. However, none of these strains have been investigated as part of two diarrheal manifestation groups for their growth and metabolic capabilities as has been done in this study. All *C. jejuni* strains were grown under microaerophilic conditions (5% O_2_, 10% CO_2_, 85% N_2_) and maintained on Mueller-Hinton agar plates supplemented with 5% defibrinated bovine blood (MHB; Becton, Dickinson and Company, Difco) for 48 hours at 37°C. Strains for assays that required liquid culture were grown in Mueller-Hinton (MH; Becton, Dickinson and Company Difco) broth with constant shaking (100 rpm) for 16-18 hours at 37°C under microaerophilic conditions.

**Table 1.** *Campylobacter jejuni* strains utilized in the study and associated diarrheal manifestation.

| Strain | Source | Clonal Complex | ST | Year of Isolation | Diarrheal Manifestation |
| --- | --- | --- | --- | --- | --- |
| RM1221 | Chicken | ST-354 | ST354 | 2005 | Watery |
| S3 | Chicken | ST-354 | ST354 | 2007 | Watery |
| D42a | Chicken | ST-21 | ST21 | 2007 | Watery |
| A14a | Chicken | ST-607 | ST1212 | 2007 | Watery |
| A13a | Chicken | ST-607 | ST1212 | 2007 | Watery |
| NCTC11168 | Clinical | ST-21 | ST43 | 1977 | Bloody and/or Inflammatory |
| 81-176 | Clinical | ST-42 | ST604 | 1985 | Bloody and/or Inflammatory |
| M129 | Clinical | ST-353 | ST353 | 1989 | Bloody and/or Inflammatory |
| A9a | Chicken | ST-460 | ST2827 | 2007 | Bloody and/or Inflammatory |
| D33a | Chicken | ST-42 | ST459 | 2007 | Bloody and/or Inflammatory |

### Growth Curves

Growth curves in MH broth were conducted for each of the *C. jejuni* strains. Overnight cultures were grown with shaking (100 rpm) at 37°C under microaerophilic conditions, OD_600_ measured, and then diluted to a final OD_600_ of 0.001 (∼1 x 10^6^ CFU/ml) with MH broth. Next, 300 µl aliquots of the diluted inoculum were added in triplicate to wells in a sterile 96-well polystyrene plate (Fisher Scientific), and the plate incubated at 37°C with under microaerophilic conditions for 1 hour before being sealed tightly with an optical plate film cover (Applied Biosystems) to prevent evaporation during incubation. The sealed plate was then placed in a SpectraMax M2 plate reader at either 37°C or 42°C, to investigate body temperature differences between humans (37°C) and chickens (42°C; the primary *C. jejuni* reservoir), with continuous shaking (100 rpm) for 36 hours with OD_600_ readings recorded every 15 minutes. Generation times for each strain were calculated using readings from the logarithmic phase of growth. Three biological replicates were performed for all 10 *C. jejuni* strains with two technical replicates (n = 3).

### Carbon Utilization Microarray

Phenotype carbon utilization microarrays were conducted as described previously by Line et al [47]. Briefly, phenotype microarrays (PMs) were conducted using the PM1 and PM2A plates (Biolog Inc., CA, USA) for variation in carbon utilization sources between strains. Each strain of *C. jejuni* was grown on MHB agar for 12 – 16 hours at 37°C under microaerophilic conditions. The bacteria were collected from the MHB agar plate using an inoculating loop, resuspended in 17 ml of IF-0a medium (Biolog Inc., CA, USA), and then diluted to a final OD_600_ of 0.8 in IF-0a medium. Next, the inoculum suspension was mixed with IF-0a, a phenotype microarray additive solution containing tetrazolium violet (Redox dye D), according to manufacturer’s instructions. Then 100 µl of the inoculum suspension was added to each well of either the PM1 or PM2A plates. The plates were incubated at 37°C under microaerophilic conditions for 48 hours before taking OD_550_ measurements in a microplate reader (SpectraMax M2) and visually recording a color change from clear to violet in each well on a scale of 0-3. A negative control plate for both types of plates was also incubated in the absence of bacteria to determine false positives. Each strain was screened a single time with both the PM1 and PM2A carbon utilization plates to serve as an initial screening method to indicate potential carbon sources of interest to study further for the strains as described below (n = 1).

### L- Fucose, L- Glutamine, and Mucin Utilization Assays

The phenotype microarray initial screening results indicated carbon utilization differences between the diarrheal manifestation groups for both L-fucose and L-glutamine, thus we focused on these carbon sources for additional studies. Studies on L-glutamine were done at both 37°C and 42°C, while L-fucose utilization assays were done at only 37°C. Mucin utilization at 37°C was added as an additional assay as it has been shown to be important to *Campylobacter* pathogenesis and L-fucose is a component of mucus. L-fucose, L-glutamine, and mucin utilization assays were adapted from Stahl et al [42]. Briefly, overnight MH broth cultures of the different *C. jejuni* strains were grown with shaking (100 rpm) then diluted to a final OD_600_ of 0.01 with minimum essential medium (MEM; Gibco) without glutamine or phenol red and supplemented with 20 μM FeSO_4_. The media was also supplemented with either no supplement, 25 mM L-fucose (Thermo Scientific), 0.01 mg/ml bovine submaxillary gland mucin (Millipore) or 20 mM L-glutamine (Alfa Aesar) depending on the assay. Next, 300 μl aliquots of the diluted inoculum with or without the appropriate supplement were inoculated in triplicate into a sterile 96-well plate, incubated at 37°C under microaerophilic conditions for 1 hour, then sealed tightly with an optical plate film cover (Applied Biosystems) to prevent evaporation during incubation. The sealed plate was then placed in a SpectraMax M2 plate reader at either 37°C or 42°C (for L-fucose and L-glutamine) with continuous shaking (100 rpm) for 36 hours with OD_600_ reading recorded every 15 minutes. The plate included a blank control well containing 300 µl of uninoculated medium. Results were recorded as the maximum OD_600_ obtained for the culture during 36 hours of growth minus the blank media OD_600_[39]. Each strain was tested in three biological replicates with three technical replicates for each of the different carbon utilization assays with either L-fucose, mucin or L-glutamine or without supplements (n = 3). There was no growth without any L-fucose, mucin or L-glutamine present during any of the assays.

### Motility Assay (Soft Agar Swarming)

Motility assays were conducted based on assays described previously by Reuter et al [48] with the following modifications: (1) using an OD_600_ of 1.0; (2) checking at time points: 24 hours and 48 hours; (3) using a 10 µl inoculation into the agar; and (4) incubating the agar plates at either 37°C or 42°C. Briefly, each *C. jejuni* strain was grown overnight in MH broth, the OD_600_ measured, and the culture diluted in MH broth to a final OD_600_ of 1.0. Mueller Hinton (MH) soft agar plates (Mueller Hinton broth supplemented with 0.4% agar) were inoculated with 10 µl of the diluted inoculum directly into the center of the plate by stabbing into the agar. Inoculated plates were incubated at either 37°C or 42°C under microaerophilic conditions for 48 hours. The radius of growth was measured after 24 hours and 48 hours of incubation using a digital calibrator starting from the edge of the inoculum spot to the edge of growth. Each strain was inoculated onto two separate plates for two technical replicates, and each assay was conducted in three biologically independent assays (n = 3).

### Biofilm Assay

Biofilm assays were performed as previously described by Reeser et al[49] except for the following modification: (1) strains were inoculated at an OD_600_ of 0.25. Briefly, *C. jejuni* strains were grown overnight in MH broth with shaking (100 rpm) at 37°C under microaerophilic conditions, the OD_600_ was measured, and the culture diluted with MH broth to a final OD_600_ of 0.25. Then 100 µl of diluted inoculum were then added to the corresponding well of a sterile 24-well polystyrene plate (Fisher Scientific, Los Angeles, CA, USA) containing 1 ml of MH broth. The plate was then statically incubated at 37°C under microaerophilic conditions for 72 hours. After 72 hours incubation, all liquid media was carefully removed from each well by pipetting to avoid disrupting the pellicle, and then the plate was dried at 55°C for 30 minutes. The pellicle in each well was then stained by adding 1 ml of 0.1% crystal violet (100 mg of crystal violet powder (Acros Organics), 8 ml of sterile double distilled water (ddH_2_O), and 2 ml 100% methanol (Fisher Bioreagents)) and incubating at room temperature for 5 minutes. Next, each well was washed twice with 1 ml of sterile ddH_2_O to remove any excess crystal violet, the plate was dried at 55°C for 15 minutes and then decolorized by adding 1 ml of decolorizer solution (80% ethanol (Fisher Bioreagents) and 20% acetone solution (Fisher Bioreagents)). Finally, 100 µl of the decolorized solution for each strain was transferred to a corresponding well of a 96-well polystyrene plate, and the absorbance at 570 nm was measured using a SpectraMax M2 microplate reader. Each strain was tested in two technical replicates during three biologically independent assays (n = 3).

### Statistical Analysis

Statistical analyses were performed using GraphPad Prism (v10.4.1) for comparisons between diarrheal manifestations or RStudio (v2022.07.1) for comparisons between strains. All data were tested for normality (Shapiro–Wilk test) and equal variance (Levene’s test). For comparisons between strains associated with watery and bloody/inflammatory diarrhea (n = 15 per group), the non-parametric Mann–Whitney U test was applied. Statistical significance was defined as p ≤ 0.05. For comparisons across multiple strains or conditions, non-parametric Kruskal–Wallis tests with Dunn’s post hoc test were used. Compact Letter Display (CLD) notation was used to visualize the results of the pairwise comparisons. Strains that do not share a letter are significantly different from each other (p < 0.05), while strains sharing the same letter are not significantly different. Statistical significance was defined as p ≤ 0.05. Each assay included a minimum of three independent biological replicates per strain (n = 3), with technical replicates as noted. Data are presented as mean ± SEM unless otherwise stated.

## Results

### Watery diarrhea-associated C. jejuni strains grow faster than bloody/inflammatory associated strains at 37°C

To evaluate whether there are differences in growth dynamics between strains associated with different diarrheal manifestations, we compared the growth kinetics of five watery diarrhea-associated and five bloody/inflammatory-associated *C. jejuni* strains at two host-relevant temperatures: 37°C (human) and 42°C (chicken). Growth was monitored over 36 hours using OD measurements taken every 15 minutes. Strain to strain differences were observed widely among both 37°C and 42°C. Strains M129 and NCTC11168 grew significantly slower at 37°C than at 42°C (p < 0.05; **Figure 1A**).

**Figure 1.**
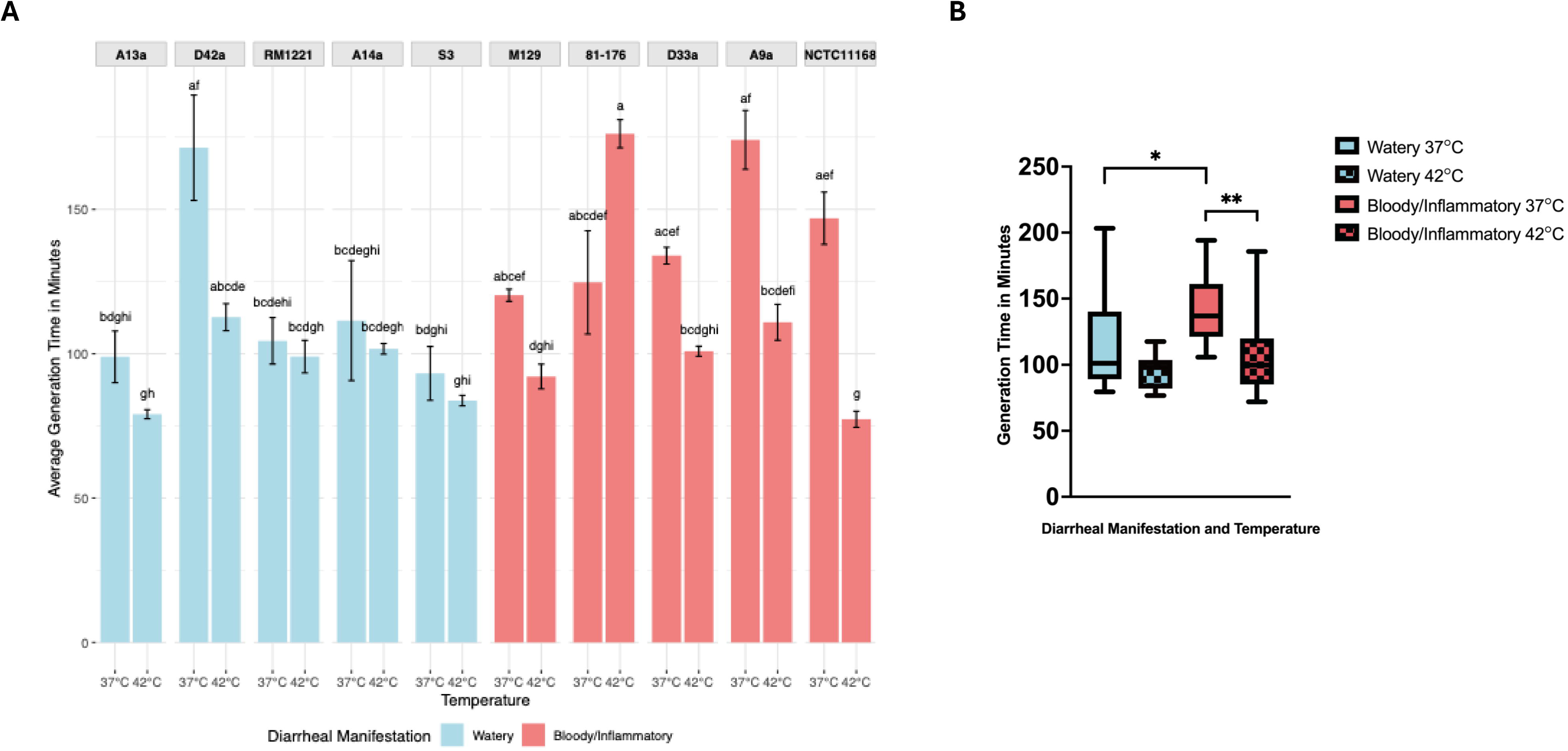
Generation time of *C. jejuni* strains at 37°C and 42°C. (A) Generation Time in minutes at 37°C and 42°C for all *C. jejuni* strains. Average generation time calculated from logarithmic phase from three biologically independent assays with three technical replicates at both 37°C and 42°C (n = 3). Compact letter displays (CLDs) indicate statistically significant differences between groups (p < 0.05) as determined by Kruskal-Wallis test followed by Dunn’s post hoc test. Bars labeled with different letters indicate statistically significant differences (p < 0.05) between those strains and/or temperatures, whereas shared letters denote groups that are not significantly different from each other. (B) Generation time in minutes of *C. jejuni* strains grouped by diarrheal manifestation (n = 15). There was a significant difference between bloody/inflammatory and watery diarrhea associated strains at 37°C (*, p = 0.018) but not at 42°C (p = 0.38). There was a significant difference between 37°C and 42°C among Bloody/inflammatory diarrhea associated strains (**, p = 0.0057), but not among watery diarrhea associated strains (p = 0.12).

At both temperatures, watery diarrhea associated strains entered logarithmic growth phase earlier and reached higher OD values compared to bloody/inflammatory strains (**Supplementary Figure 1A–D**). One watery strain (D42a) displayed a slower growth profile, similar to that of bloody/inflammatory strains. Generation times calculated from logarithmic growth phase found significantly faster generation times for watery diarrhea associated strains compared to bloody/inflammatory associated strains at 37°C but not at 42°C (37°C watery diarrhea associated 115.8 minutes, 37°C bloody/inflammatory diarrhea associated 139.99 minutes p = 0.018; 42°C watery diarrhea associated 95.23 minutes, bloody/inflammatory diarrhea associated 42°C 111.47 minutes p = 0.38; **Figure 1B**). There was no significant difference in generation times at 37°C compared to 42°C for watery diarrhea associated strains (p = 0.12) but bloody/inflammatory diarrhea associated strains grew significantly faster at 42°C than 37°C (p = 0.0057).

### Strains differ in utilization of L-fucose and L-glutamine but not mucin

To determine whether there are nutrient utilization differences between strains associated with distinct diarrheal outcomes, we screened all ten *C. jejuni* strains for their ability to metabolize 192 carbon sources using Biolog’s PM1 and PM2A phenotype microarray plates. Several carbon sources, including L-glutamic acid, succinic acid, fumaric acid, a-hydroxy-butyric acid and D-L-malic acid were utilized by all ten *C. jejuni* strains, regardless of their diarrheal manifestation. The amino acid L-aspartic acid was utilized strongly by watery diarrhea associated strains A13a, S3 and weakly by RM1221 but was more strongly utilized by bloody/inflammatory strains M129, 81-176, NCTC11168 and D33a, as indicated by a stronger color change. There were a few carbon sources used solely by one or two watery diarrhea associated strains and not by bloody/inflammatory diarrhea associated strains and vice versa, including D-tagatose (D33a), succinnamic acid (D33a, 81-176), citric acid (NCTC11168 and 81-176) and tricarballyic acid (S3). Several carbon sources exhibited distinct utilization patterns between the two diarrheal manifestation groups (**Supplementary Figure 2**). The sugar D-arabinose was utilized by two watery diarrhea associated strains D42a and S3, but was utilized more strongly by three bloody/inflammatory diarrhea strains (M129, 81-176, NCTC11168). L-serine was utilized by only three bloody/inflammatory diarrhea associated strains (M129, 81-176 and NCTC11168) but not utilized by any watery diarrhea associated strains, while both L-glycyl-proline and glycyl-L-glutamic acid were utilized by watery diarrhea associated strains A13a, A14a, S3 but only bloody/inflammatory diarrhea associated strain A9a. Most notably, the amino acid L-glutamine and the sugar L-fucose were utilized by multiple strains in only one group: L-glutamine was used by three bloody/inflammatory strains (81-176, M129, D33a) and only one watery strain (S3), while L-fucose was used by four watery strains (A13a, A14a, D42a, S3) and just one bloody/inflammatory strain (A9a). The possible role that L-fucose and L-glutamine utilization, along with mucin, could have in colonization of the host’s intestinal tract among these *C. jejuni* strains made them desirable for further phenotypic analysis [50–52].

All five watery strains showed robust growth in the presence of L-fucose at 37°C (**Figure 2A**). The geometric mean of the OD for watery strains was 0.076 compared to 0.030 for bloody/inflammatory strains (p < 0.0001). L-fucose supported significantly greater growth in watery diarrhea-associated strains (**Figure 2B**). These results suggest that L-fucose metabolism is a consistent trait among watery diarrhea-associated strains. To assess whether differences in mucosal substrate metabolism extended beyond individual monosaccharide components of mucin, like L-fucose, we next tested the ability of each strain to grow in medium supplemented with bovine submaxillary gland mucin at 37°C. Although there was strain-level variability, no significant difference in mucin utilization was observed between watery and bloody/inflammatory associated strains (p = 0.099), (**Figure 3, Supplementary Figure 3**).

**Figure 2.**
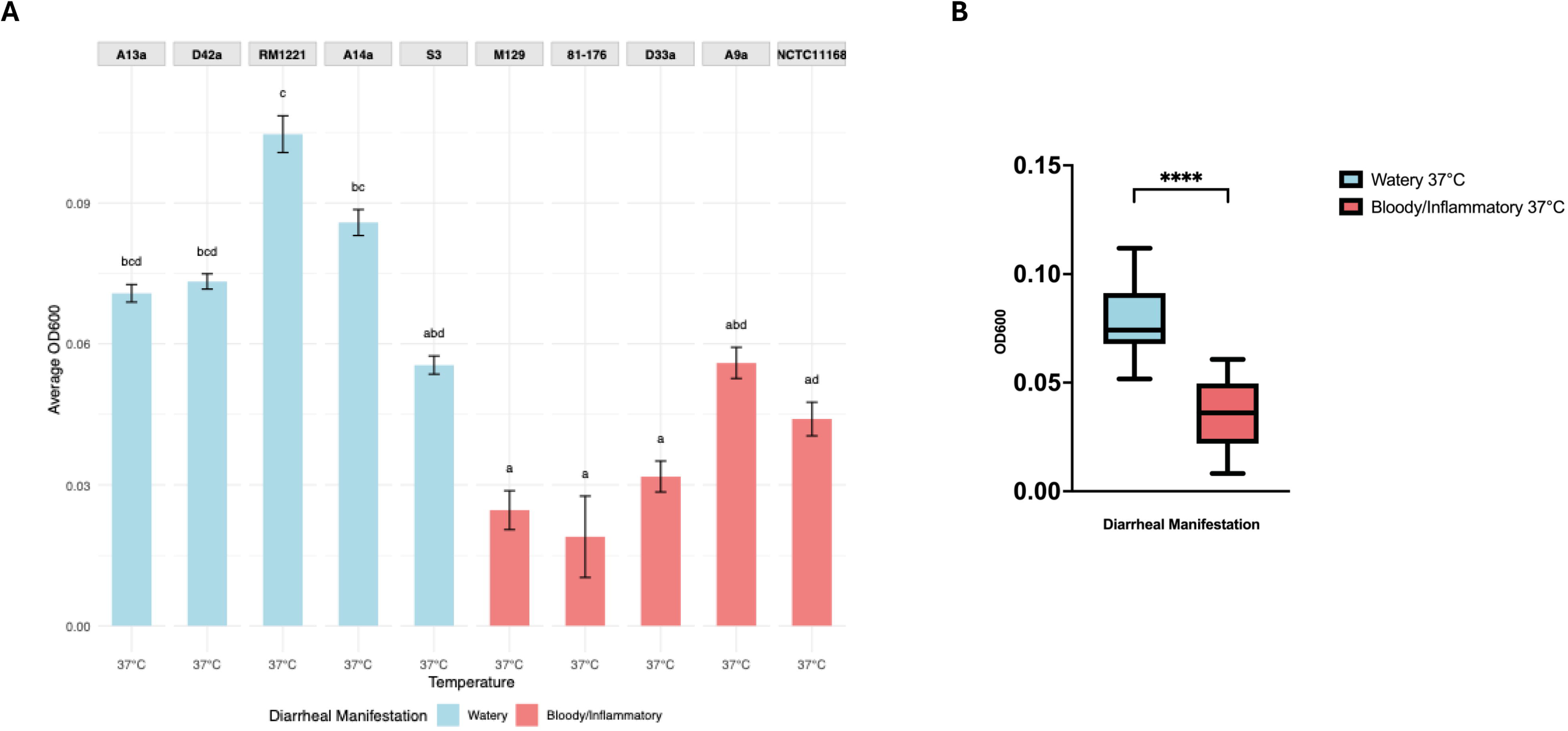
Utilization of L-fucose at 37°C. (A) L-fucose utilization at 37°C between *C. jejuni* strains. Compact letter displays (CLDs) indicate statistically significant differences between groups (p < 0.05) as determined by Kruskal-Wallis test followed by Dunn’s post hoc test (n = 3). Bars labeled with different letters indicate statistically significant differences (p < 0.05) between those strains whereas shared letters denote groups that are not significantly different from each other. (B) L-fucose utilization at 37°C between *C. jejuni* strains grouped by diarrheal manifestation. There was a significant difference between the growth of watery diarrhea and bloody/inflammatory associated strains when grown in minimal media containing L-fucose at 37°C (****, p < 0.0001). The average highest OD_600_ was calculated from three biologically independent assays with three technical replicates (n = 15).

**Figure 3.**
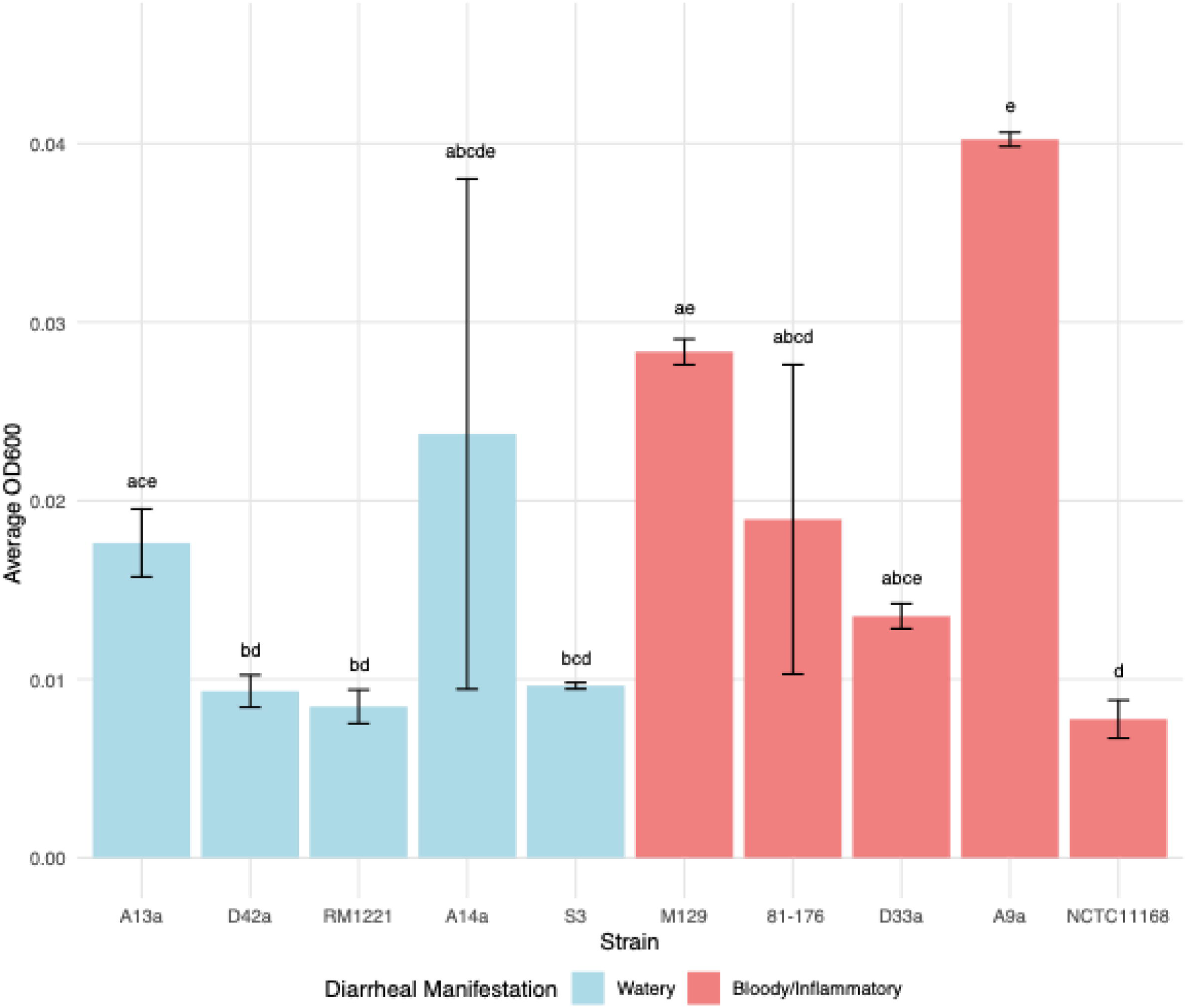
Utilization of mucin at 37°C. Bovine submucosally gland mucin utilization at 37°C by *C. jejuni* strains. Compact letter displays (CLDs) indicate statistically significant differences between groups (p < 0.05) as determined by Kruskal-Wallis test followed by Dunn’s post hoc test. Bars labeled with different letters indicate statistically significant difference (p < 0.05) between those strains, whereas shared letters denote groups that are not significantly different from each other. The average highest OD_600_ was calculated from three biologically independent assays with three technical replicates (n = 3).

L-glutamine utilization supported higher levels of growth among all strains compared to L-fucose, with several significant differences between different strains at 37°C or 42°C (**Figure 4A**). Among all strains tested, only watery diarrhea-associated strain A13a showed significantly higher growth at 42°C compared to 37°C, while bloody/inflammatory diarrhea-associated strains 81-176 and A9a showed increased growth at 37°C compared to 42°C. However, when grouped by diarrheal manifestation, bloody/inflammatory associated strains exhibited enhanced growth at 37°C in L-glutamine-supplemented medium compared to watery diarrhea associated strains (p = 0.0005) (**Figure 4B**). There was no difference in growth potential at 42°C between watery and bloody/inflammatory strains, (p = 0.68), suggesting a temperature or host dependent role for glutamine utilization among *C. jejuni* strains. Together, these results demonstrate that carbon source utilization differs consistently between strain and diarrheal manifestation groups, and these traits may be modulated by host temperature. The enhanced L-fucose metabolism of watery diarrhea associated strains may facilitate rapid growth and colonization in human hosts, while the slower growth and utilization of L-glutamine by bloody/inflammatory diarrhea associated strains may contribute to persistence within the human host.

**Figure 4.**
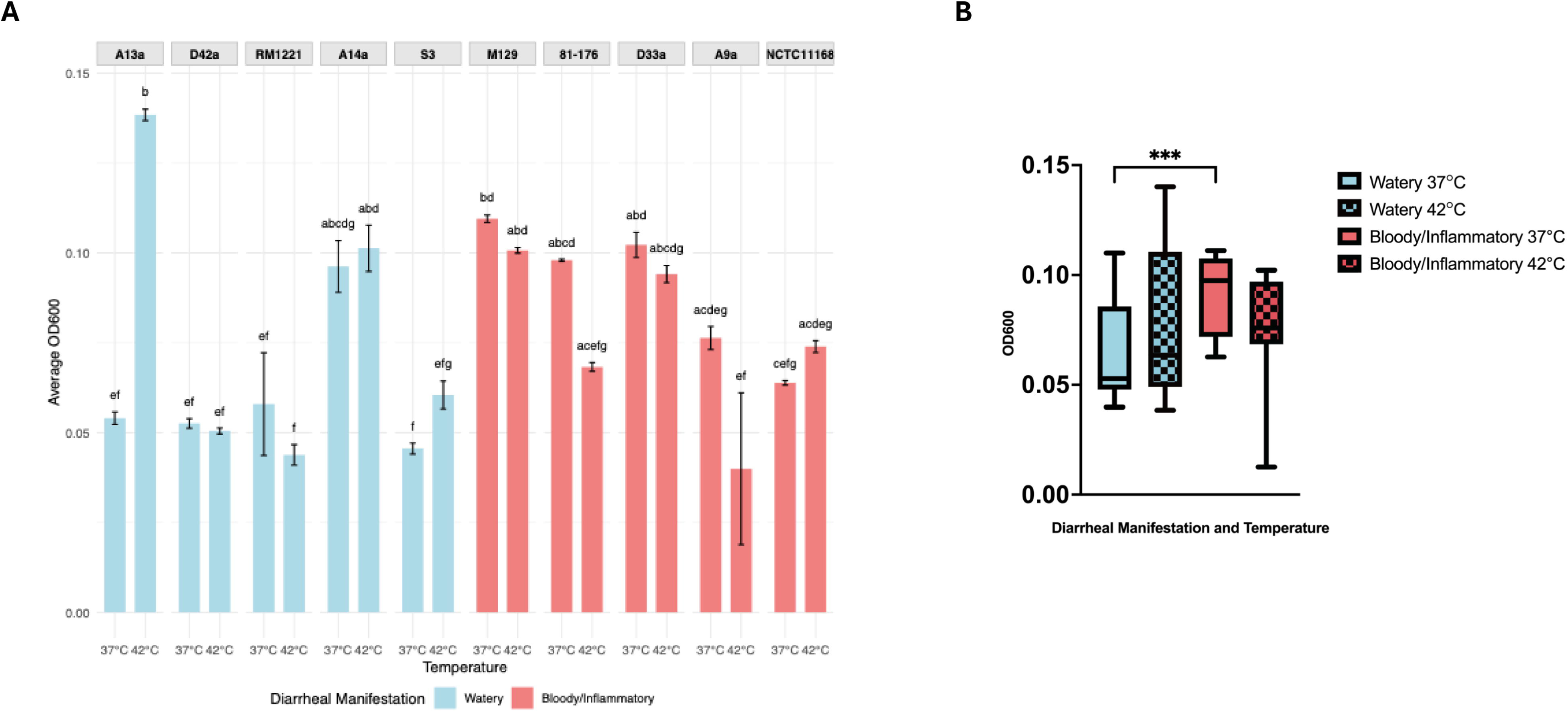
Utilization of L-glutamine at 37°C and 42°C. (A) Average highest OD_600_ L-glutamine utilization at 37°C and 42°C across all *C. jejuni* strains. Compact letter displays (CLDs) indicate statistically significant differences between groups (p < 0.05) as determined by Kruskal-Wallis test followed by Dunn’s post hoc test (n = 3). Bars labeled with different letters indicate statistically significant difference (p < 0.05) between those strains and/or temperatures, whereas shared letters denote groups that are not significantly different from each other. (B) Average highest OD_600_ L-glutamine utilization at 37°C and 42°C across all *C. jejuni* strains grouped by diarrheal manifestation. There was a significant difference between the growth of watery diarrhea and bloody/inflammatory associated strains when grown at 37°C in minimal media containing L-glutamine (***, p = 0.0005) but not at 42°C (p = 0.68). The average highest OD_600_ was calculated from three biologically independent assays with three technical replicates (n = 15).

### Motility varies widely among C. jejuni strains but does not distinguish diarrheal phenotype

Motility is critically important for *C. jejuni* colonization and pathogenesis, aiding in both adhesion as well as invasion of host cells [53–55]. To assess whether motility differs between strains associated with either watery or bloody/inflammatory diarrhea, we performed swarming motility assays in soft agar at both 37°C and 42°C. A wide range of motility phenotypes were observed among *C. jejuni* strains: watery diarrhea associated strains D42a and RM1221, and bloody/inflammatory diarrhea associated strains NCTC11168, D33a, and 81-176 were highly motile (>10 mm radius), while watery diarrhea associated strain S3 and bloody/inflammatory diarrhea associated strain A9a were weakly motile or non-motile (<10 mm) (**Figure 5**). Strains A13a and A14a both had higher (but not significantly different) levels of motility at 42°C. No significant difference in motility was observed between strains associated with the different diarrheal manifestations at either temperature: 37°C (p = 0.33) or 42°C (p = 0.64; **Supplementary Figure 4**). There was no significant difference in motility at different temperatures for either the watery (p = 0.35) or bloody/inflammatory (p = 0.24) diarrhea associated strains.

**Figure 5.**
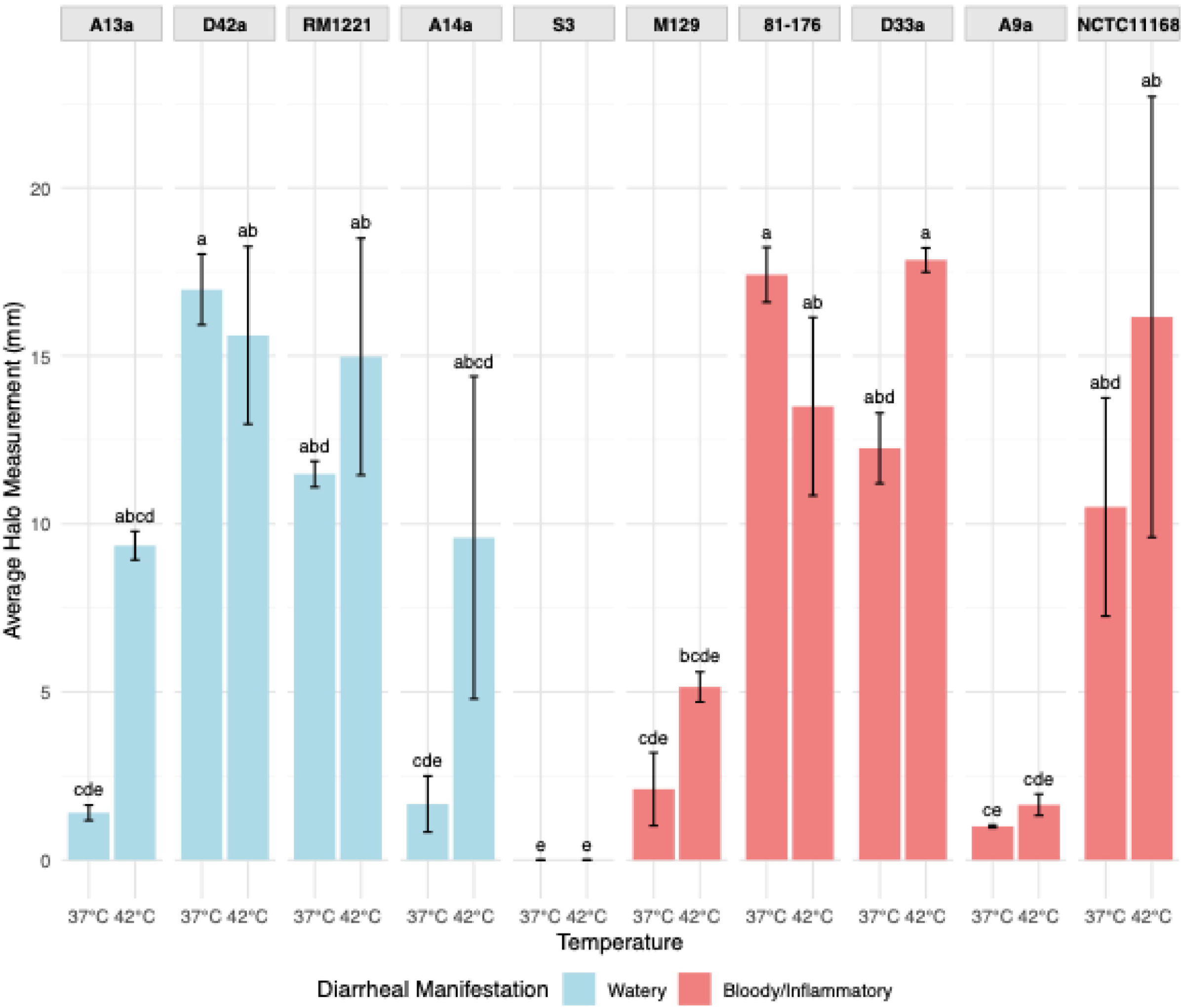
Motility at 37°C and 42°C. Average motility of *C. jejuni* strains after 48-hour incubation at both 37°C and 42°C. Motility was measured as the radius of the halo of growth in millimeters after 48-hour incubation. Average motility was calculated from three biologically independent assays with two technical replicates (n = 3). Compact letter displays (CLDs) indicate statistically significant differences between groups (p < 0.05) as determined by Kruskal-Wallis test followed by Dunn’s post hoc test (n = 3). Bars labeled with different letters indicate statistically significant difference (p < 0.05) between those strains and/or temperatures, whereas shared letters denote groups that are not significantly different from each other.

### Biofilm formation does not differ by diarrheal phenotype

Biofilm formation contributes to the persistence, and transmission of *C. jejuni* [56]. Therefore, to assess the potential differences in these traits among our *C. jejuni* strains, we quantified biofilm formation at 37°C using crystal violet staining. Strong biofilm producers (A > 0.30) included watery diarrhea associated strains A13a and D42a and bloody/inflammatory diarrhea associated strain M129. Medium biofilm formation (A between 0.20–0.30) was observed in watery diarrhea associated strains A14a and S3 and bloody/inflammatory diarrhea associated strains 81-176 and NCTC11168, while poor biofilm formation (A < 0.20) was noted for watery diarrhea associated strain RM1221 and bloody/inflammatory diarrhea associated strains A9a and D33a (**Figure 6A**). Although strains varied in their individual capacities, biofilm formation at 37°C was not found to significantly differ between diarrheal manifestation groups (p = 0.061; **Figure 6B**).

**Figure 6.**
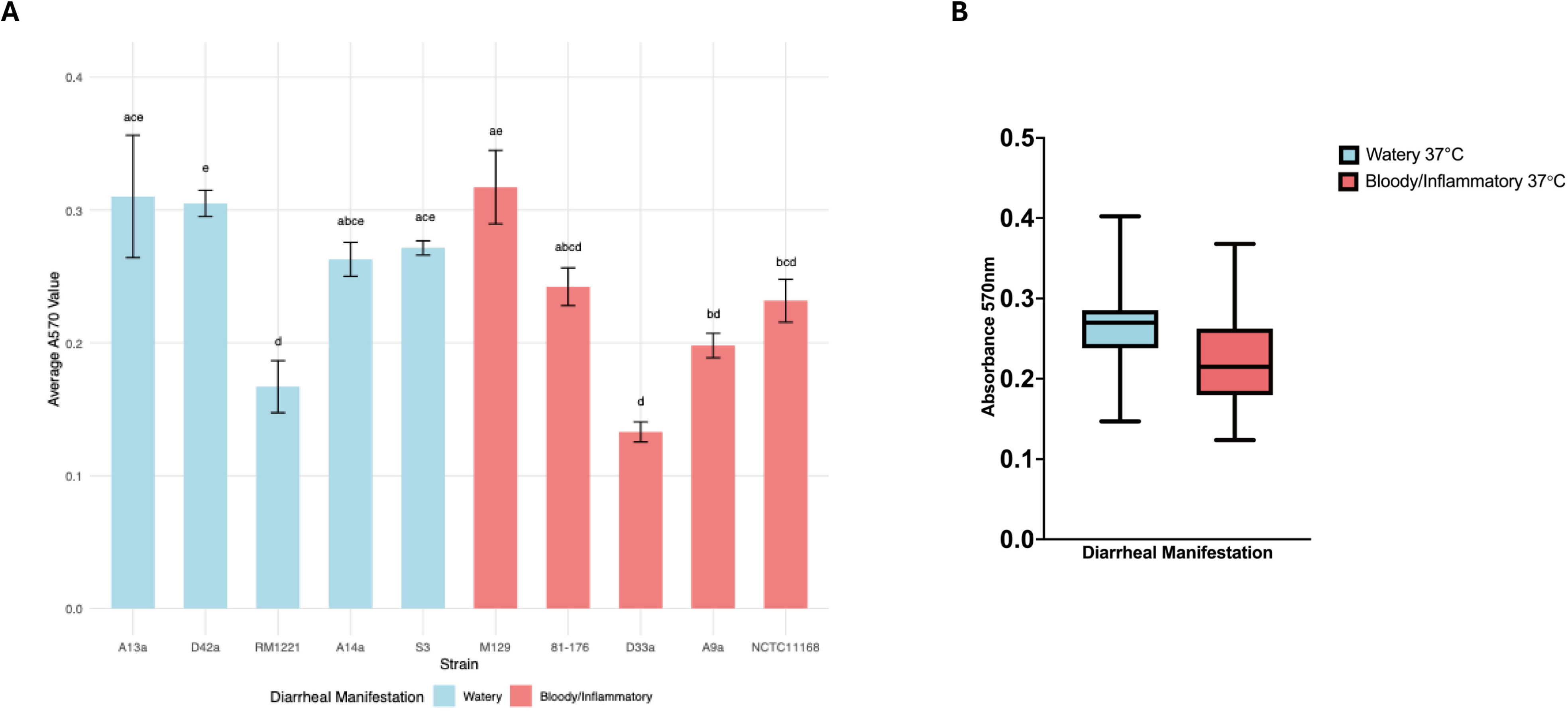
Biofilm production at 37°C. (A) Formation of biofilm after 72 hours at 37°C among *C. jejuni* strains. Biofilm formation was quantified using spectrometer readings at 570 nm following a 72-hour incubation at 37°C. Compact letter displays (CLDs) indicate statistically significant differences between groups (p < 0.05) as determined by Kruskal-Wallis test followed by Dunn’s post hoc test. Bars labeled with different letters indicate statistically significant difference (p < 0.05) between those strains, whereas shared letters denote groups that are not significantly different from each other. Average biofilm formation was determined from three biologically independent assays at 37°C with two technical replicates (n = 3) (B) Average biofilm production of watery diarrhea and bloody/inflammatory associated strains after 72 hours at 37°C. Average biofilm production at 37°C was calculated from three biologically independent assays with two technical replicates (n = 15).

## Discussion

This study investigated the variation between strains of *C. jejuni* with defined diarrheal phenotypes in the neonatal piglet model, we assessed growth rates, carbon source utilization, motility, and biofilm formation at human and chicken body temperatures. We observed that watery diarrhea-associated *C. jejuni* strains grow faster than bloody/inflammatory associated strains at 37°C. Here, *C. jejuni* strains showed significant strain to strain variation, including differences in the utilization of L-fucose, L-glutamine, generation times, and but did not display significantly different abilities to form biofilms or to utilize bovine submaxillary gland mucin when grouped by diarrheal manifestation, a finding that was not surprising based on *C. jejuni’s* limited ability to metabolize carbohydrates [35]. The uniqueness in carbon utilization between diarrheal manifestation groups may be attributed to numerous factors including host adaptation, evolutionary advantages or preferred niches.

We observed very little to no growth of any of the *C. jejuni* strains in the presence of mucin, suggesting an inability to break it down on their own. It is known that *C. jejuni* growth is enhanced when co-cultured with commensal *Bacteroides vulgatus*, a response suggested to be due to the breakdown of mucin by *B. vulgatus* [57]. There are several types of bacteria that are known mucin degraders that have been identified in humans including *Bifidobacterium* [58, 59], *Bacteroides* [59] and *Akkermansia muciniphila* [60]. Mucin degradation by *A. muciniphila* is known to support intestinal barrier function [61] and provide physiological benefits that prevent disease [61–63]. The composition of mucin degraders in chickens is less well defined, but *Bacteroides* has been found to be one of the most predominating genera of bacteria in the chicken intestinal microbiome [62, 64]. The growth of these commensal bacteria is supported by the highly glycosylated mucins within the intestinal mucus [65], during which active fructosidases are secreted [66, 67]. *C. jejuni* lacks the ability to release L-fucose from glycans and instead is known to utilize free fucose from mucin degraders such as *B. vulgatus* [57].

The mucus layer of the intestinal tract is where *C. jejuni* encounters L-fucose, the only known carbohydrate chemoattractant for *C. jejuni* [68, 69]. In the human gut, 80% of the total dry weight of intestinal mucins is made up of *O*-linked glycans, while L-fucose is a key terminal glycan the exact percentage is not known due to the diversity in glycosylation between individuals [70–72]. However, in chickens L-fucose is suggested to comprise an estimated 7% of the *O*-glycans in the intestinal tract [73]. The *fuc* locus for L-fucose catabolism has been identified in >50% of sequenced *C. jejuni* isolates [74] and has been observed in a high percentage of *C. jejuni* isolates from humans [75]. Here we found that strains associated with watery diarrhea exhibited enhanced utilization of L-fucose at 37°C. These watery diarrhea associated strains also grew significantly faster overall at 37°C than those linked to bloody/inflammatory diarrhea. While these differences may be observational in nature, we hypothesize that this combination of rapid growth and efficient metabolism may support rapid early colonization, helping to contribute to the acute nature of watery diarrhea. Reduced invasiveness by watery diarrhea associated strains may also be a key factor to the observed diarrheal manifestation, as the bacteria would remain within the mucus layer of the gut where it rapidly grows and is eventually expelled [76].

Chemotaxis towards L-fucose in the intestinal lumen through the upregulation of L-fucose dehydrogenase, FucX, is suggested to contribute to its clearance while non-L-fucose metabolizing *C. jejuni* strains move towards the intestinal mucosa to establish disease [77]. These findings would support the hypothesis that watery diarrhea associated strains utilize L-fucose in the mucus layer where they are maintained further away from the intestinal epithelial cells while bloody/inflammatory diarrhea associated strains move to establish themselves at the enterocyte surface. Indeed, we have recently found that bloody/inflammatory diarrhea associated *C. jejuni* strains are significantly more invasive than watery diarrhea associated strains [76], a finding that may suggest that a higher level of virulence among bloody/inflammatory diarrhea associated strains could be a key contributor to the diarrhea manifestation differences. However, in contrast, L-fucose utilization has also been observed *in vitro* to enhance invasion of Caco-2 cells and significantly increase fibronectin binding efficacy in strain NCTC11168 [44]. Another study found similar results with invasion of Caco-2 increased with the addition of 10 mM L-fucose under microaerophilic conditions using *C. jejuni* strain 108 [67]. The same study observed growth enhancement of *C. jejuni* strain 108 in the presence of L-fucose, a finding similar to ours, suggesting that watery diarrhea associated strains may alter their invasiveness if tested under the same conditions. In addition, we found that the increased growth potential of watery diarrhea associated strains was maintained at both host-relevant temperatures, suggesting adaptation to colonization across multiple hosts with enhanced growth in humans. The ability to utilize L-fucose provided a colonization advantage *in vivo* in piglets but had no effect on the colonization of chickens when inoculated at a low dose [42]. Since piglets are a model for human disease, whereas chickens are the natural reservoir for *C. jejuni*, this suggests L-fucose utilization at 37°C (human body temperature) versus 42°C (chicken body temperature) may only be relevant to human disease. Other factors including virulence factors that have not yet been elucidated may also strongly contribute to these observed growth and metabolic differences in strains associated with one diarrheal manifestation compared to the other.

Bloody/inflammatory associated diarrhea strains grew more slowly particularly at 37°C, a factor that potentially contributes to prolonged colonization and disease development. Slower growth may allow for slower establishment of an infection, which allows for delayed immune recognition of the pathogen until the infection is more established, potentially resulting in a more damaging infection. Initial Biolog screening found that some strains preferentially utilized L-glutamine, we observed a significant increase in growth potential in the presence of L-glutamine by bloody/inflammatory associated strains compared to watery diarrhea associated strains at 37°C but not at 42°C. In humans, glutamine is known to support epithelial barrier function and modulate host immune responses, and its depletion may promote tissue damage and inflammation [78–80]. Because L-glutamine is a key fuel source for enterocytes and supports tight junction integrity [78, 81], *C. jejuni* utilization of L-glutamine may provide a metabolic advantage and also actively disrupt enterocyte homeostasis. In the context of inflammation, where glutamine demand is heightened due to increased epithelial turnover and immune cell activation, its depletion could contribute to impaired repair to tissue damage as well as reduced immune function [82]. Other studies have found that *C. jejuni* strains have considerable variation in their ability to utilize L-glutamine, with 31% of strains [40] containing the enzyme gamma-glutamyl transpepidase (GGT) necessary for glutamine and glutathione metabolism. One study found a higher percentage of *C. jejuni* strains from human isolates (37.93%) to contain the *ggt* gene compared to 14.29% among strains from non-human sources [51], suggesting an advantage for these strains in the human host. Furthermore, a study of 166 *C. jejuni* strains from Finnish patients found an association between bloody diarrhea and GGT production [83], while another study found purified *C. jejuni* GGT impaired intestinal cell and lymphocyte proliferation [84]. The complicated and variable response by *C. jejuni* strains containing *ggt* suggests that individual strains may have complex metabolic requirements or can compensate with alternative pathways not yet characterized. Overall, the metabolic diversity observed among strains highlights the importance of considering host-specific and strain-specific adaptations when investigating L-glutamine utilization among *C. jejuni* strains, especially among those associated with bloody/inflammatory diarrhea.

This study found that motility differences were not restricted to the different diarrheal manifestation groups. Motility was variable at the strain level at either 37°C or 42°C. All strains were motile except strain S3, which was originally able to colonize chickens [85, 86], but minimal laboratory passages rendered it defective for colonization [87]. Interestingly, motility is a critical factor associated with colonization/pathogenicity [88–91]. Previously, similar results were demonstrated for non-motile *C. jejuni* strain CS that caused few pathological changes in the piglet model, but grew to a greater maximum cell density in MH medium compared to a genotypically similar but motile strain [89]. Watery diarrhea associated strains with limited to no motility, such as S3, may be able to overcome the mucus layer via the more rapid growth that we observed in this study, but additional studies are needed to confirm this hypothesis. Our results demonstrate that *C. jejuni* strain motility does not have a role in the type of diarrheal manifestation associated with the strain.

Biofilm formation is known to differ among *C. jejuni* strains [92] and has been suggested to promote survival, transmission and colonization of hosts [93, 94]. The present study revealed that there was no significant difference in biofilm formation among watery or bloody/inflammatory diarrhea associated strains at 37°C, thus demonstrating that factors other than biofilm formation during human colonization are important to the type of diarrheal manifestation produced by the strain. In addition to temperature, biofilm formation by bacteria can be triggered by various environmental stressors such as nutrient availability, varying oxygen tensions and osmotic changes [95–97], for example, one study showed that *C. jejuni* biofilm formation was increased under low-oxygen tensions as well as in minimal nutrient medium [49]. When nutrients are available, such as L-fucose, it is known that biofilm formation is reduced, and growth is enhanced *in vitro* [74]. Future studies should investigate the ability of *C. jejuni* strains associated with watery diarrhea to form biofilms when L-fucose is provided in the medium, particularly given their enhanced growth with L-fucose.

It is important to note that our study revealed substantial phenotypic variability within each diarrheal manifestation group, and there are likely numerous *C. jejuni* associated factors and host factors associated with the different diarrheal manifestations. In fact, it is not unexpected to see strain to strain variability given the known genomic and functional diversity of *C. jejuni*. However, this strain to strain variability reinforces the need to study multiple strains to identify group-level trends. Within the watery diarrhea group, strain D42a had a longer generation time at 37°C than the other watery associated strains. A13a showed significantly higher growth at 42°C in the presence of L-glutamine. Among bloody/inflammatory strains, NCTC11168, and M129 had slower generation times at 37°C than at 42°C. 81-176, D33a and NCTC-11168 showed similar levels of motility, but A9a and M129 were only weakly motile. The presence of these outliers highlights the spectrum of metabolic and growth capabilities within each diarrheal manifestation group, and further supports that these diarrheal manifestations are probably due to numerous different virulence and host factors. As mentioned previously, that bloody/inflammatory diarrhea associated strains may simply have enhanced virulence factors compared to watery diarrhea associated strains, an explanation supported by other work by our lab [76]. Additional work is necessary to determine if growth rate and metabolic traits along with various key virulence attributes are true contributors to these two diarrheal manifestations in human disease.

Another consideration is that strain classification in this study was based on diarrheal outcomes observed in the neonatal piglet model, rather than direct human clinical data. While this may limit immediate generalizability, the piglet model is the only experimental systems that reliably reproduces both watery and bloody/inflammatory diarrheal manifestations caused by *C. jejuni.* It provides a controlled environment to directly associate specific bacterial traits with clinical outcomes, eliminating many of the confounding variables inherent in human observational studies. Additionally, clinical strains are likely biased towards bloody/inflammatory diarrhea associated strains, as people suffering from the less severe watery diarrhea are not likely to seek medical attention. Nonetheless, validation of these findings using well-characterized human isolates with known clinical presentations will be important for confirming their broader relevance. Furthermore, there are a wide range of confounding factors that exist that could further contribute to the diarrheal manifestations, even within the neonatal piglet model. The piglet model used with these ten *C. jejuni* strains in prior studies were conventional not gnotobiotic piglets, allowed for slight variation in the gut microbiome. It remains to be elucidated if these *Campylobacter* strains may differ in their abilities to outcompete and/or interact with the gut microbiome in piglets and/or human during disease production.

Despite these limitations, our findings provide initial evidence that the diarrheal outcome produced during a *C. jejuni* infection may be at least partially shaped by fundamental differences in the growth and metabolism of the infecting strain. Strains associated with watery diarrhea appear optimized for rapid expansion and mucus-derived sugar utilization, while those linked to bloody/inflammatory diarrhea are slower growers, utilizing L-glutamine for enhanced growth at 37°C. These strategies may reflect differing host adaptation trajectories and suggest that metabolic profiling could provide new insight into the pathogenic potential of *C. jejuni* lineages.

## Contributors

**JB:** Conceptualization, methodology, experimentation, formal analysis, writing – original draft, and review & editing. **KKC:** Conceptualization, methodology, resources, funding acquisition, project administration, supervision, writing, and review & editing. **CP:** Writing, and review & editing. **BP:** Writing, and review & editing.

## Acknowledgments

The authors thank all members of the Cooper laboratory for providing critical technical assistance in sample processing and other aspects of the study. The authors thank Bryan Roxas for assistance with Supplementary Figure 2 and thank Dr. Jared Smith for his contributions to editing the manuscript.

