## Supplementary Figures for "Growth Rates and Metabolic Traits Differ by Diarrheal Manifestation in *Campylobacter jejuni* Strains"

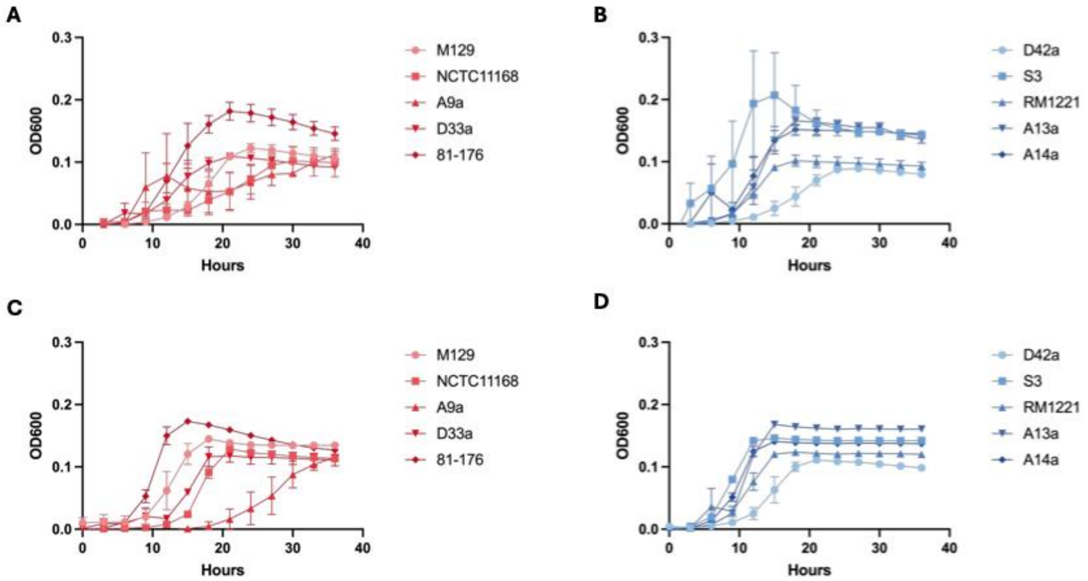


**Supplementary Figure 1. Growth curves of *C. jejuni* strains in MH broth at 37°C and 42°C.** (A) Growth curve performed over 36 hours for bloody/inflammatory diarrhea associated strains at 37°C. (B) Growth curve performed over 36 hour for watery diarrhea associated strains at 37°C. (C) Growth curve performed over 36 hours for bloody/inflammatory diarrhea associated strains at 42°C. (D) Growth curve performed over 36 hours for watery diarrhea associated strains at 42°C. For each of the growth curves, the timepoint represents the average $OD_{600}$ of each strain between three biologically independent assays performed with three technical replicates (n = 3), and data was collected every hour for 36 hours and displayed in 3 hour increments.

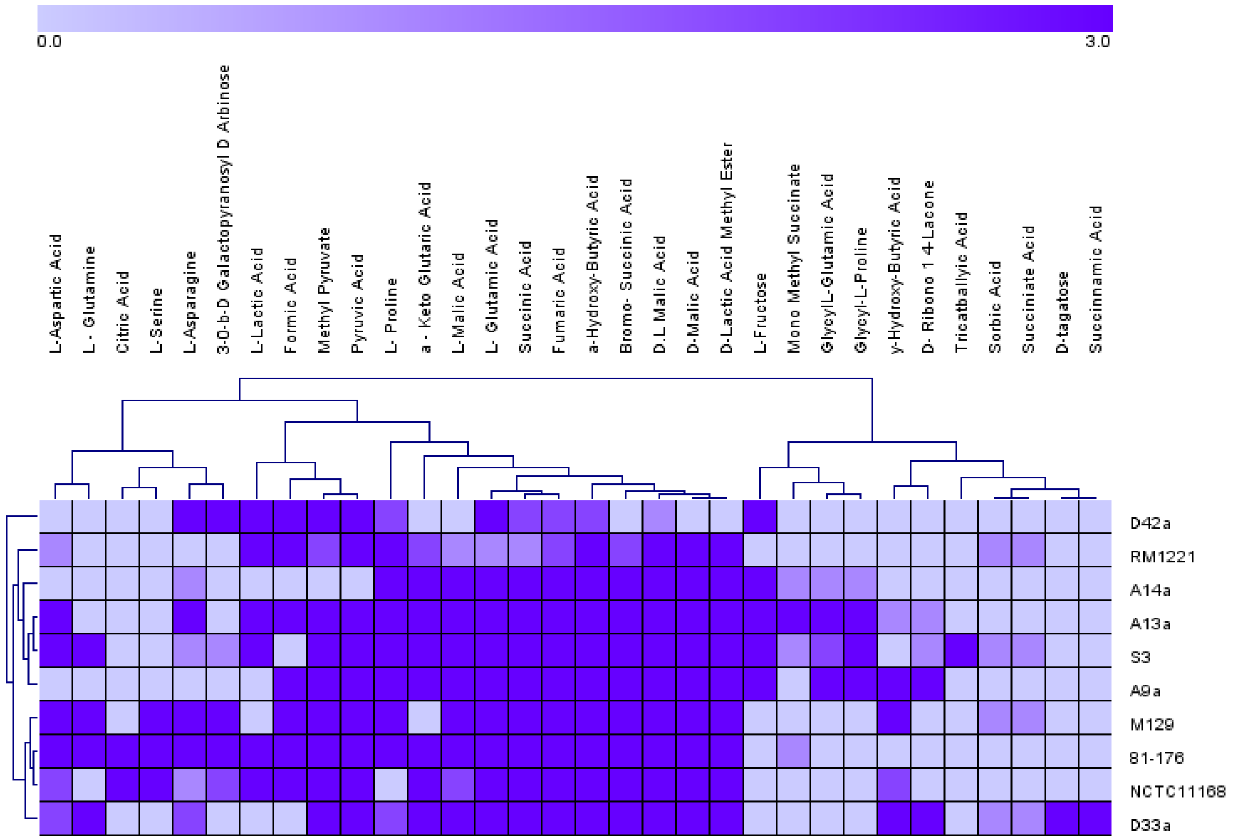


**Supplementary Figure 2. Utilization of carbon sources from BIOLOG PM1 and PM2 among *C. jejuni* strains.** Heat map showing all carbon sources from BIOLOG PM1 and PM2 plates that appeared positive, indicated by the change in the clear media a purple color. The intensity of the purple color was ranked on a scale of 0 to 3, with 0 being no color and 3 being an intense purple color. BIOLOG PM1 and PM2 plates were performed once per strain (n = 1).

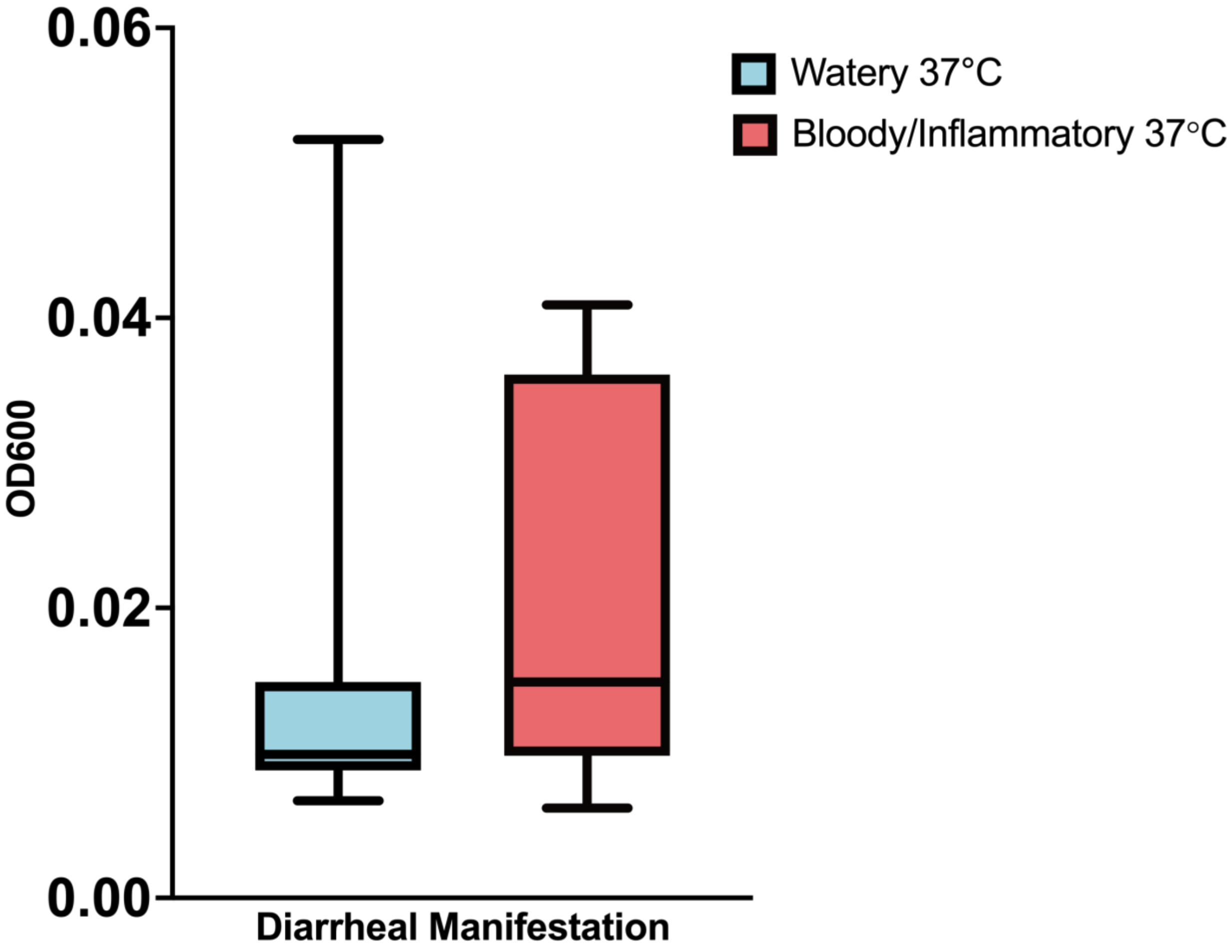


**Supplementary Figure 3. Utilization of bovine submaxillary gland mucin at 37°C among *C. jejuni* strains grouped by diarrheal manifestation.** Average highest $OD_{600}$ of bovine submaxillary gland mucin utilization at 37°C and 42°C across all *C. jejuni* strains grouped by diarrheal manifestation (n = 3). There was no significant difference between the growth of watery and bloody/inflammatory diarrhea associated strains when grown in minimal media containing bovine submaxillary gland mucin (p = 0.099).

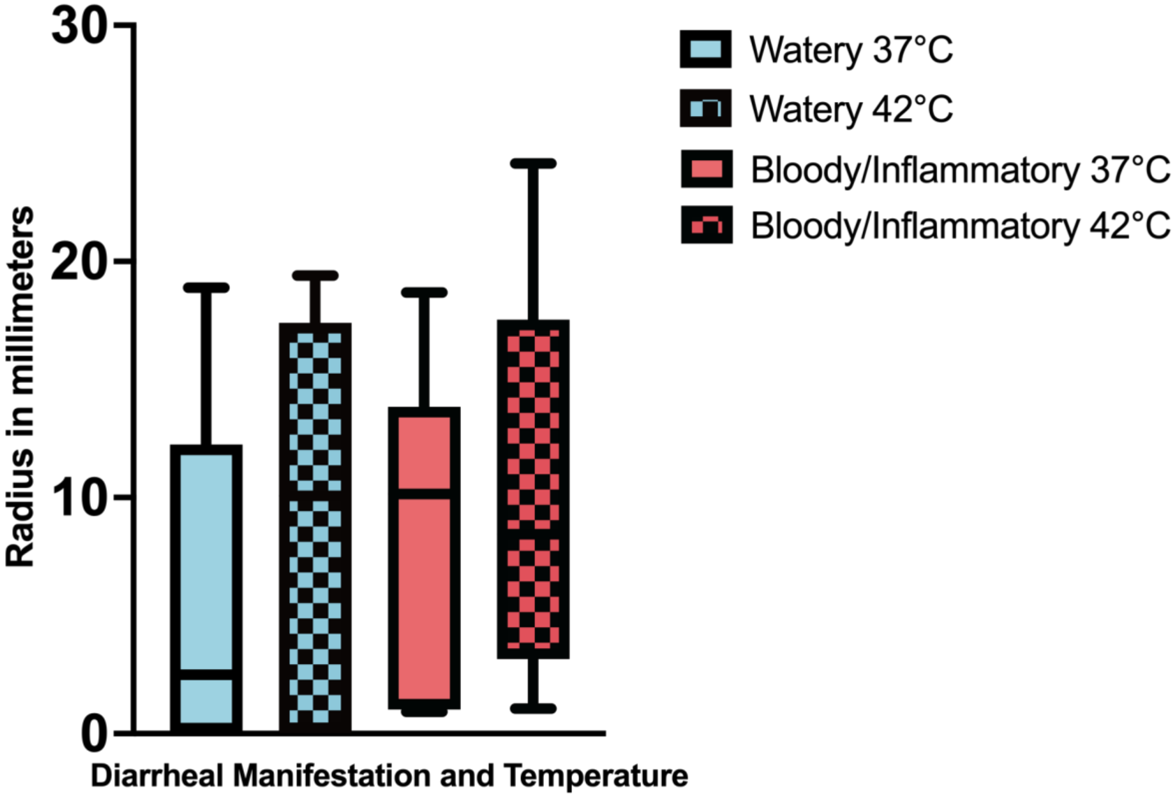


**Supplementary Figure 4. Average motility of *C. jejuni* strains grouped by diarrheal manifestation at 37°C and 42°C.** Motility was measured as the radius of the halo of growth in millimeters 48 hours. There was no significant difference in motility between watery and bloody/inflammatory diarrhea associated strains at both 37°C ($p = 0.33$) and 42°C ($p = 0.64$). There was no significant difference between the motility of bloody/inflammatory diarrhea associated strains ($p = 0.24$) or watery diarrhea associated strains at 37°C and 42°C ($p = 0.35$). Average motility at 37°C and 42°C was calculated from three biologically independent assays with two technical replicates ($n = 3$).
